# Silicon Micropillar-Enhanced CRISPR Biosensor for Rapid and Sensitive Detection of Drug-Resistant Bacteria

**DOI:** 10.64898/2025.12.15.694528

**Authors:** Ruonan Peng, Zhenxuan Yuan, Demis D. John, Biljana Stamenic, Fatt Foong, Bhaskar Sharma, FNU Yuqing, Chujing Zheng, Jacob Waitkus, Ke Du

## Abstract

The growing threat of antibiotic-resistant pathogens, such as methicillin-resistant Staphylococcus aureus (MRSA), underscores the urgent need for rapid, sensitive, and field-deployable diagnostic technologies. Here, we present a silicon micropillar-enhanced CRISPR biosensor that integrates high-aspect-ratio microstructures with a one-pot RPA/CRISPR-Cas12a assay for ultrasensitive and specific detection of MRSA. Micropillar arrays with fixed diameters and varying heights (100 µm, 300 µm, and 500 µm) were fabricated via deep reactive ion etching and functionalized for surface probe immobilization. Suboptimal crRNA design was employed to modify Cas12a activation kinetics, enabling declined trans-cleavage and enhanced end-point signal accumulation. The 500 µm micropillar configuration demonstrated a tenfold improvement in sensitivity compared to the 100 µm array, with a limit of detection reaching 10^3^ CFU mL^-1^. The platform also showed high specificity against non-target bacterial strains. These findings highlight the potential of combining microstructured chips with one-pot CRISPR diagnostics to advance next-generation point-of-care tools for infectious disease monitoring.

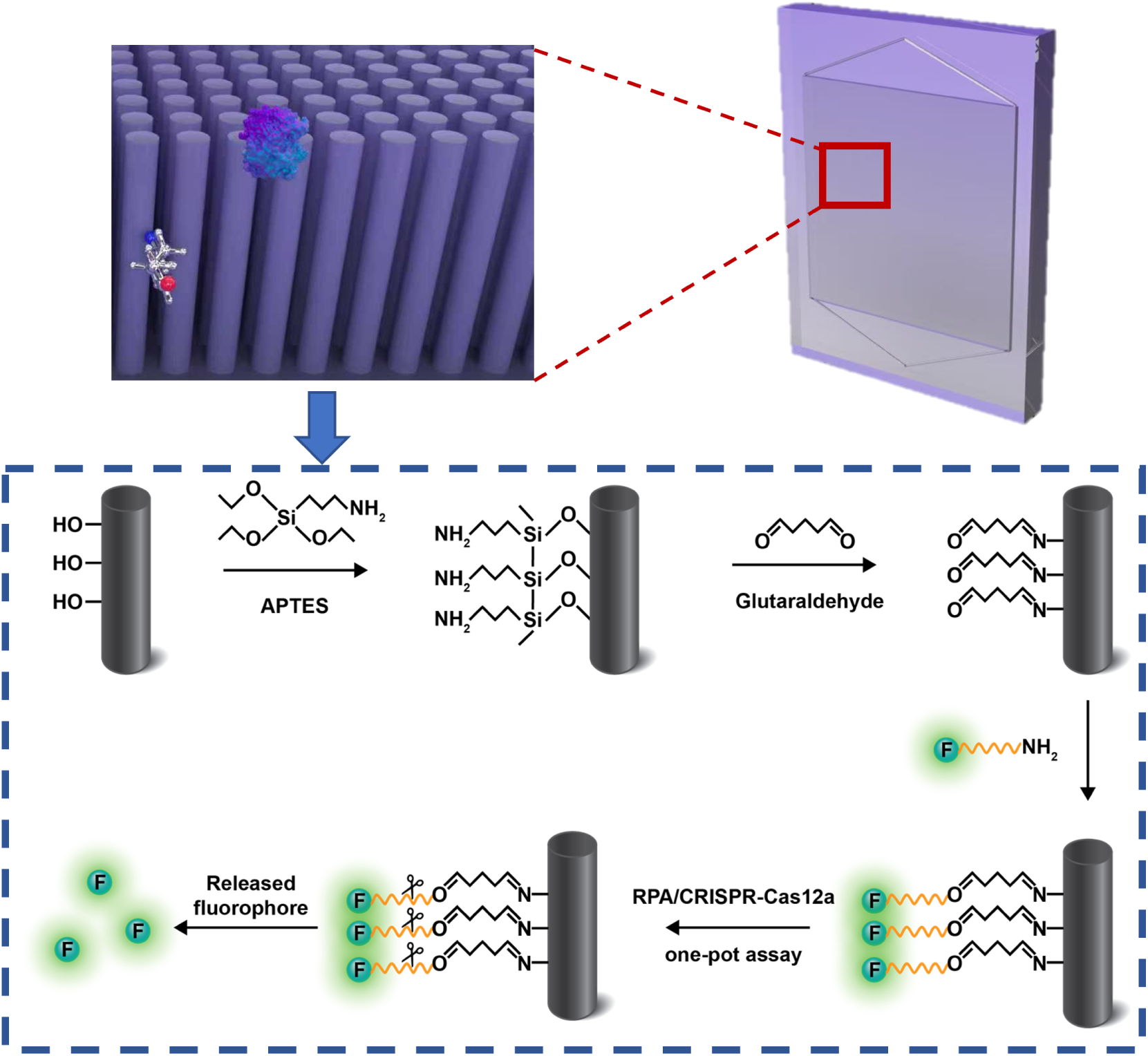

## 2 Introduction

The global rise of antibiotic resistance represents an urgent threat to public health, undermining the effectiveness of standard treatments and increasing the incidence of severe and persistent infections (Salam et al., 2023). In the United States alone, antibiotic-resistant infections are responsible for an estimated 23,000 deaths annually and contribute to over $20 billion in additional healthcare costs (Habboush and Guzman, 2023). Among these drug-resistance pathogens, methicillin-resistant *Staphylococcus aureus* (MRSA) is particularly concerning due to its resistance to beta-lactam antibiotics and widespread occurrence in both healthcare and community settings (Gajdács, 2019). MRSA can cause a variety of infections, including skin and soft tissue infections (Kaushik et al., 2024), pneumonia (Lemelle et al., 2025), bloodstream infections (Boattini et al., 2025), and surgical site infections (Ahmann et al., 2025), many of which can become life-threatening if not promptly diagnosed and treated (Mandal et al., 2024). Traditional detection methods, such as bacterial culture and PCR-based assays, often suffer from limitations including long turnaround times, susceptibility to contamination (Haley et al., 2024), reliance on thermal cycling and specialized laboratory equipment (He et al., 2023). These challenges have driven growing interest in developing point-of-care (POC) diagnostic platforms that are rapid, sensitive, and easy to deploy, particularly in resource-limited environments (Heidt et al., 2020).

Clustered regularly interspaced short palindromic repeats (CRISPR) and associated (Cas) nucleases, particularly Cas12 and Cas13 nucleases, have emerged as powerful tools for *in vitro* molecular diagnostics due to their programmability (Jiao et al., 2024), high specificity (Mann and Pitts, 2022), and collateral cleavage activity (Palaz et al., 2021). Among them, Cas12a has attracted significant attention for DNA detection. It uses a single CRISPR RNA (crRNA) to recognize double-stranded DNA targets adjacent to a specific Protospacer Adjacent Motif (PAM), typically 5’-TTTV-3’ (where V = A, C, or G) (Ai et al., 2025). Upon target recognition, Cas12a becomes activation and initiates two types of nuclease activity: *cis*-cleavage, which cuts the target DNA at the distal end of PAM (Lin et al., 2025), and *trans*-cleavage, which enables indiscriminate cleavage of surrounding single-stranded DNA (ssDNA) (Zhou et al., 2024). This *trans*-cleavage activity forms the foundation of signal amplification in CRISPR-based biosensors, in which cleavage of fluorophore–quencher (FQ) probes releases fluorescence upon activation (Zhang et al., 2021; Peng et al., 2023). Additionally, Cas12a operates efficiently at 37 °C, making it more compatible with POC applications than thermal cycling-based PCR methods (Gootenberg et al., 2018).

To further improve detection sensitivity, the CRISPR-Cas12a system is often coupled with isothermal amplification techniques such as recombinase polymerase amplification (RPA) (Peng et al., 2025). RPA amplifies target DNA at a constant temperature through the combined action of recombinases, single-stranded DNA-binding proteins (SSBs), and strand-displacing DNA polymerases. In this process, recombinases facilitate the pairing of primers with complementary target sequences, SSBs stabilize the displaced DNA strands, and DNA polymerases extend the primers in the presence of deoxynucleotide triphosphates (dNTPs), resulting in rapid and efficient amplification (Daher et al., 2016). While the combination of RPA and CRISPR enables rapid and sensitive detection, conventional two-step workflows, where amplification and detection occur in separate stages, require manual reagent transfers, increasing procedural complexity and the risk of cross-contamination (Yin et al., 2020). In attempts to merge the two reactions into a single one-pot assay introduces new challenges. Cas12a may be activated prematurely, leading to early cleavage of RPA amplicons before sufficient amplification occurs. This reduces the overall template concentration and limits signal output (Lu et al., 2022; Wang et al., 2025). Moreover, FQ probes inherently exhibit background fluorescence due to incomplete quenching. Excess probe concentrations can increase this baseline signal, reducing signal-to-noise ratio and ultimately limiting detection sensitivity (Hass et al., 2020).

To overcome these limitations, microfabricated platforms with micro- and nanostructured features have played a pivotal role in advancing biosensing and biological applications by enabling enhanced sensitivity (Siavashy et al., 2024), reduced sample volumes (Salva et al., 2021; Zhu et al., 2021), and overall device miniaturization (Fu et al., 2024). Among these, micropillar arrays have attracted considerable interest due to their ability to significantly increase the available surface area for probe immobilization, thereby enhancing target capture efficiency and reaction kinetics (Chen et al., 2020). However, the fabrication of high-aspect-ratio microstructures with consistent fidelity presents significant technical challenges. Achieving reproducible architectures requires precise control over cleanroom-based processes such as photolithography, dry etching, and process timing. Even slight deviations in these parameters can result in non-uniform features, structural defects, or mechanical instability, which complicates large-scale implementation. Despite these challenges, several recent studies have implemented micropillar-based diagnostics for nucleic acid detection utilizing fluorescence (Bao et al., 2024), electrochemical (Movilli et al., 2020), or colorimetric readouts (Geissler et al., 2020). Yet, few of these systems incorporate CRISPR with upstream isothermal amplification in a one-pot format. Integration of one-pot RPA/CRISPR assays into microstructured platforms remains underexplored, primarily due to challenges such as synchronizing amplification and detection kinetics within confined microenvironments, achieving stable probe immobilization, and maintaining reagent compatibility. Overcoming these hurdles would unlock the full potential of CRISPR diagnostics on chip-based systems, enabling portable, sensitive, and field-deployable biosensors.

In this study, we present a silicon micropillar-enhanced CRISPR biosensor designed for the rapid, sensitive, and specific detection of MRSA. The platform leverages high-density surface probe immobilization combined with a one-pot RPA/CRISPR-Cas12a reaction to streamline assay workflows and minimize contamination risk. To enhance the probe binding capacity and improve signal generation, we fabricated high-aspect-ratio silicon micropillars using deep reactive ion etching, with structures featuring fixed diameters and heights of 100, 300, and 500 µm. We demonstrate that increased pillar height correlates with significantly higher probe immobilization, enabling more efficient detection. In contrast, the flat chip exhibited minimal signal difference between positive and negative samples, indicating insufficient surface area for effective probe loading and target interaction. Furthermore, by designing suboptimal crRNAs to modulate Cas12a activation kinetics, we achieved improved signal amplification and specificity. The optimized 500 µm micropillar chip demonstrated a tenfold improvement in sensitivity compared to the 100 µm configuration, highlighting the strong link between device geometry and assay performance. These findings underscore the promise of integrating microfabricated platforms with CRISPR-Cas diagnostics to advance next-generation molecular detection systems for infectious disease monitoring.

## 3 Experimental section

Details on chip fabrication, micropillar surface treatment, bacterial culture, and CRISPR-based detection are provided in the Supporting Information (SI)

## 4 Results and discussion

In this study, we used silicon micropillar-enhanced CRISPR biosensor for rapid and sensitive detection of drug-resistant bacteria. **Fig. 1a** is a photograph of the physical device, both the top and bottom sides of the chip are designed with triangular features to facilitate reagent distribution. A schematic overview of the fully assembled device is shown in **Fig. 1b**, showing the chip enclosed within the container and sealed with the cap. The silicon chip is first placed into a custom-designed container, and the container is demonstrated by filling with the red food dye. Pre-mixed RPA/CRISPR reagents are then dispensed onto the micropillar side of the chip. To minimize evaporation and prevent cross-contamination during isothermal amplification at 42 °C, the container is sealed with a soft silicone cap.

**Figure 1:**
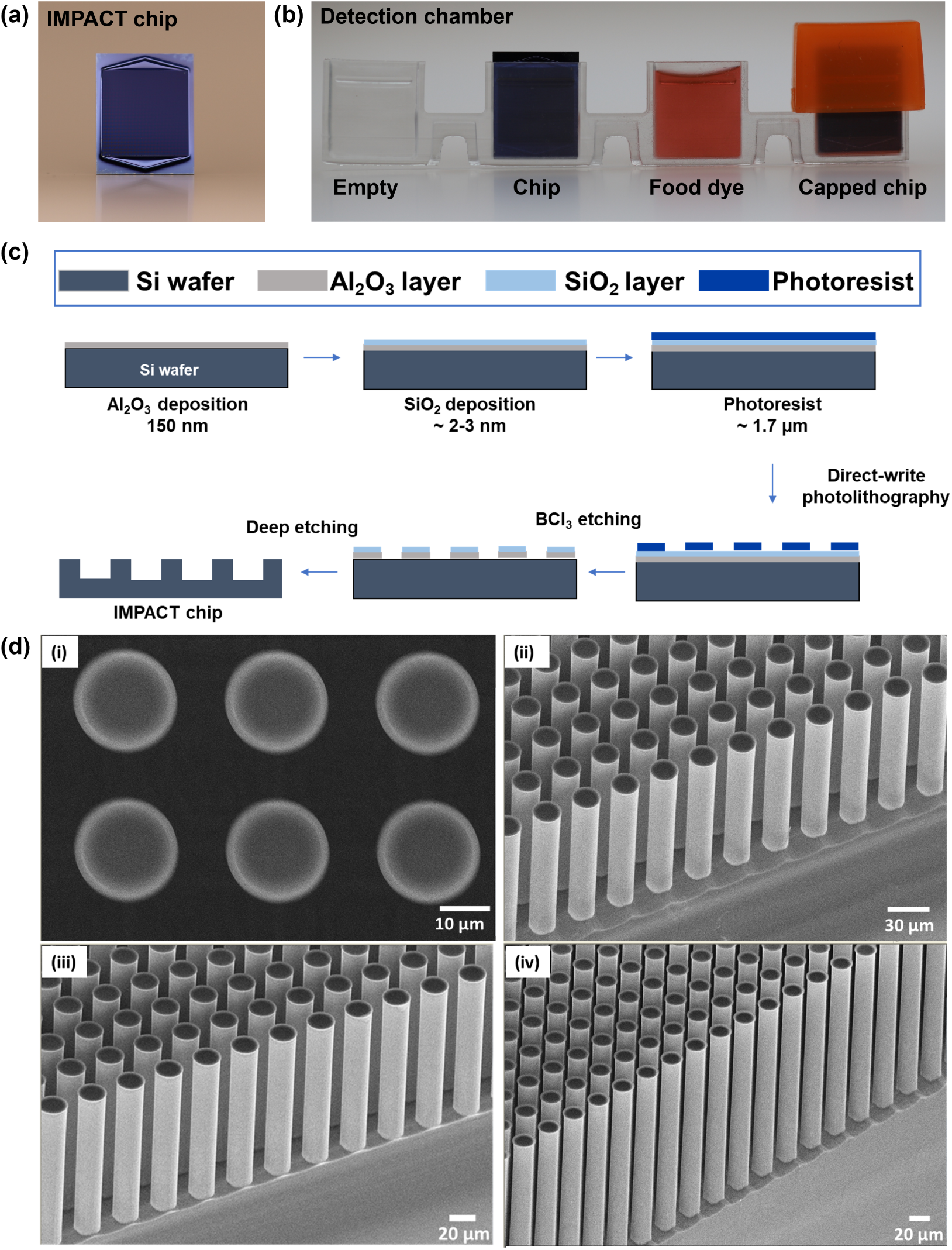
Illustration of the design, fabrication, and structural characterization of the micropillar chip. (a) Photograph of the IMPACT chip. (b) Photograph of the fabricated IMPACT chip integrated into the detection chamber. (c) Fabrication process of IMPACT chip. (d) SEM images of micropillars. (i) Top view of micropillars. (ii), (iii), and (iv) side view of micropillars with the height of 150 µm, 300 µm and 500 µm, respectively.

**Fig. 1c** presents a schematic of the microfabrication process used to construct the micropillar array. To ensure mechanical integrity after 500 µm-deep etching, *<*100*>* silicon wafers were used for high-aspect-ratio deep reactive ion etching (DRIE). Following solvent cleaning with acetone and isopropanol, a dual-layer hardmask consisting of 150 nm Al_2_O_3_ and a ∼2–3 nm SiO_2_ cap was deposited using Ion Beam-Assisted Sputter Deposition (IBD). This system was selected for its automated operation and relatively high deposition rates, making it well-suited for this process. Lithography was performed using ∼1.7 µm of nLOF2020 negative photoresist on an I-Line Heidelberg MLA150 direct-write tool. Al_2_O_3_ served as the primary hardmask due to its high etch selectivity in Bosch processes (Drost et al., 2022). However, a known challenge with Al_2_O_3_ is its susceptibility to etching in TMAH-based developers, which can cause photoresist delamination during development (Wilson et al., 2015; Ali et al., 2021). This issue was mitigated by adding a thin SiO_2_ cap layer: thick enough to protect the underlying alumina during development, yet thin enough (∼2–3 nm) to be fully removed during the subsequent BCl_3_-based ICP etch without requiring an additional step. While other deposition techniques such as ALD or sputtering are also compatible with this hardmask system, IBD was chosen for its ease of use and process reliability (Huang et al., 2024). After patterning, the oxide hardmask was etched using a BCl_3_ ICP process optimized for Al_2_O_3_, which simultaneously removed the thin SiO_2_ cap layer. The underlying silicon was etched using a modified three-step Bosch process. Process parameters were optimized via design of experiments (DOE) to achieve aspect ratios of ∼50:1. Following DRIE, the hardmask was removed, and a conformal SiO_2_ passivation layer was produced via thermal oxidation. Finally, the wafers were diced into individual chips, with special precautions, such as reduced water flow and isopropyl alcohol rinsing, taken to protect the delicate high-aspect-ratio micropillars during dicing and drying. **Fig. 1d** are SEM images of the fabricated micropillar arrays with varying heights of 100 to 500 µm. Image (i) shows a top-down view highlighting the uniform spacing and circular arrangement of the micropillars. Images (ii–iv) display tilted side views of the arrays, revealing consistent vertical alignment and well-defined cylindrical structures across different heights. These features ensure a high surface-to-volume ratio and reproducible fluid interactions critical for downstream biochemical assays.

**Fig. 2a** presents a schematic illustrating the surface functionalization process for a silicon micropillar array and its application in a surface-immobilized CRISPR-Cas12a detection assay. The process begins with plasma treatment to hydroxylate the silicon surface, introducing reactive hydroxyl groups. Then, (3-aminopropyl)triethoxysilane (APTES) is used for silanization by forming covalent siloxane bonds with the surface and introduces terminal amine (–NH_2_) groups (Vandenberg et al., 1991). The amine-functionalized surface is subsequently treated with glutaraldehyde (GA), a bifunctional aldehyde, which reacts with surface amines to form imine (Schiff base) linkages, leaving free aldehyde groups available for further conjugation (Aggarwal and Ikram, 2025). A fluorophore-labeled, amine-modified single-stranded DNA (ssDNA) probe is then covalently immobilized onto the surface via reaction with the exposed aldehyde groups (Ni et al., 2023). Upon introduction of the target DNA, a one-pot RPA/CRISPR-Cas12a reaction is initiated. The target sequence is first amplified by RPA, then recognized by the Cas12a, which is activated and exhibits nonspecific trans-cleavage activity (Li et al., 2024). This results in the cleavage of the immobilized DNA probes, releasing the fluorophores into the surrounding solution. After the chip is removed, the released fluorescence signal in the solution serves as a readout for the presence of the target sequence.

**Figure 2:**
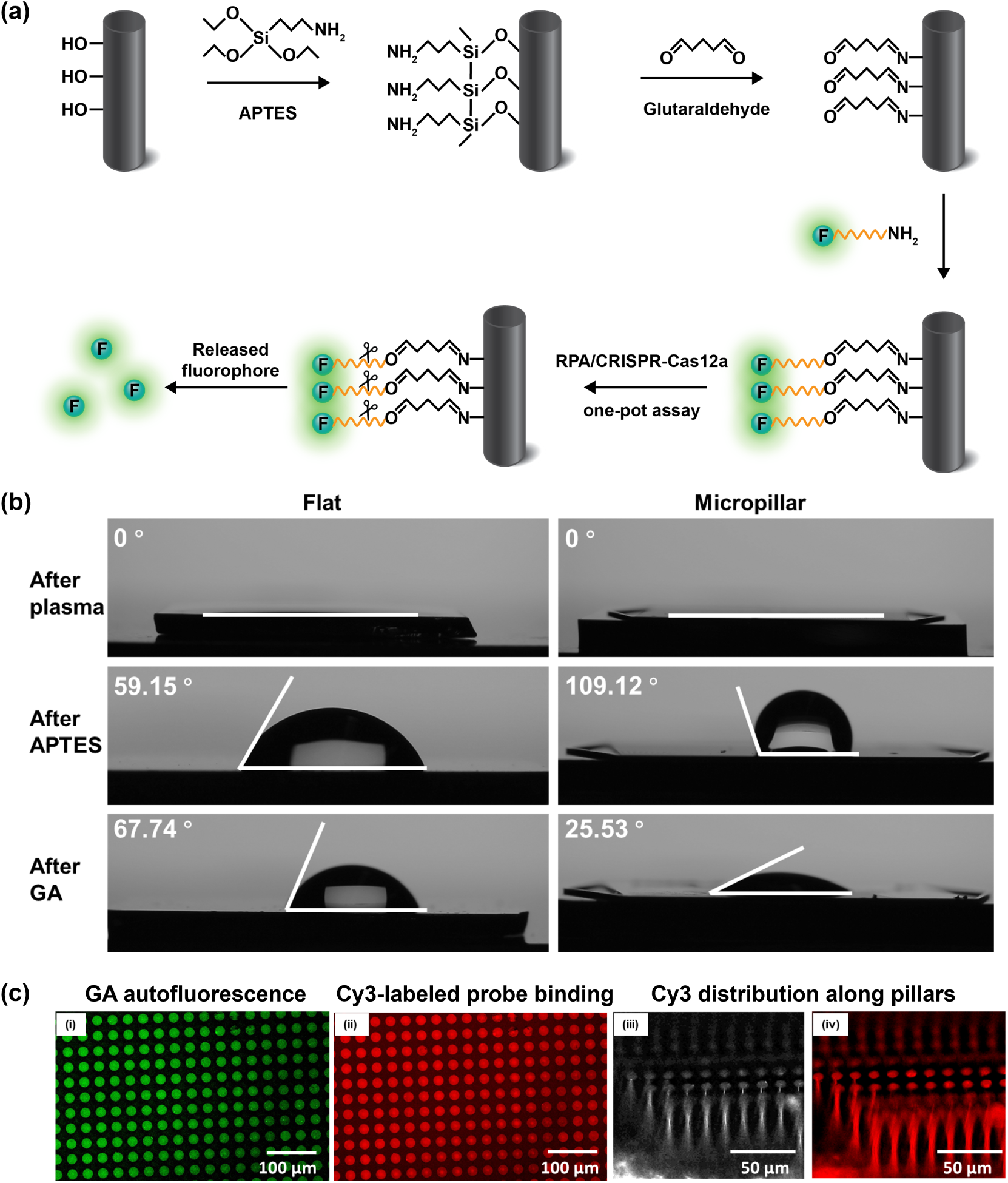
Surface treatment and characterization of the micropillars.(a) Schematic illustration of the surface functionalization process, single-stranded DNA (ssDNA) probe immobilization, and the one-pot RPA/CRISPR-Cas12a detection workflow. (b) Water contact angle measurements of flat and micropillar chips following plasma and APTES treatments. (c) Fluorescence images of 300 µm-tall micropillars after Cy3-labeled probe binding. (i) Top view under FITC filter: green fluorescence indicates GA presence. (ii) Top view under TRITC filter: red fluorescence corresponds to Cy3-labeled probes. (iii) Side view under bright field microscopy. (iv) Side view under TRITC filter showing Cy3 fluorescence distribution.

**Fig. 2b** presents the water contact angle measurements following plasma and APTES treatments for both the flat silicon chip (left) and the micropillar chip (right). Following plasma treatment, both surfaces exhibited a contact angle of 0°, indicating highly hydrophilic surfaces due to the formation of surface hydroxyl groups. After APTES treatment, the contact angle increased to 59.15° for the flat chip and 109.12° for the micropillar chip. This increase reflects the introduction of amine-functionalized alkyl chains from APTES, which impart moderate hydrophobicity. The much higher contact angle on the micropillar chip is attributed to the surface roughness amplifying the hydrophobic effect through the Cassie-Baxter wetting regime, where air pockets become trapped between the pillars, reducing liquid–solid contact (Deng et al., 2020). Following GA treatment, the flat surface exhibited a slight increase in contact angle to 67.74°, likely due to additional aldehyde functionalization contributing limited changes in surface energy. In contrast, the micropillar surface showed a substantial decrease in contact angle to 25.53°, indicating a wetting regime transition from Cassie-Baxter to Wenzel (Cwickel et al., 2017). This shift suggests that GA increased the surface energy sufficiently for water to infiltrate the microstructures, resulting in full wetting and enhanced apparent hydrophilicity. **Fig. 2c** presents fluorescence microscopy images that confirm successful surface functionalization and probe immobilization on 300 µm-tall micropillars. The images were acquired after incubation with Cy3-labeled ssDNA probes. In **Fig. 2c**, Panel (i) shows a top-view image obtained using a FITC filter, where green fluorescence indicates the presence of glutaraldehyde. This signal arises from the intrinsic fluorescence of GA due to Schiff base formation, confirming that the aldehyde layer remains intact on the surface (Song et al., 2018). Panel (ii) is a top-view image captured with a TRITC filter, showing red fluorescence corresponding to the Cy3 label. The strong, uniform fluorescence signal confirms successful immobilization of the Cy3-labeled probes onto the functionalized surface. Panel (iii) shows a side-view image under bright-field illumination, which reveals the physical structure and vertical alignment of the micropillars. Panel (iv) displays a side-view TRITC fluorescence image, demonstrating a consistent distribution of Cy3 fluorescence along the entire height of the pillars. This uniform signal suggests effective and homogeneous probe attachment throughout the 3D microstructure, validating the robustness of the surface chemistry and the accessibility of the reactive groups even at the base of the pillars.

**Fig. 3a** displays the sequences of the target DNA and corresponding crRNAs. The target site is located within the mecA gene, which encodes methicillin resistance by encoding penicillin-binding protein 2a (PBP2a). This protein exhibits low affinity for *β*-lactam antibiotics, allowing bacterial cell wall synthesis to continue despite antibiotic treatment and thereby conferring methicillin resistance (Fishovitz et al., 2014). A key element in CRISPR-Cas12a targeting is the Protospacer Adjacent Motif (PAM), which dictates where Cas12a binds and cleaves. The canonical PAM recognized by Cas12a is 5’-TTTV-3’, where V represents A, C, or G (Ai et al., 2025). In this study, two crRNAs were designed for the one-pot RPA/CRISPR-Cas12a assay: crRNA1 (5’-CCTA-3’) and crRNA2 (5’-TTCT-3’) target sequences adjacent to suboptimal PAMs. **Fig. 3b** illustrates the workflow of the one-pot RPA/CRISPR-Cas12a reaction, emphasizing the interaction between target amplification and Cas12a-mediated trans-cleavage. In the presence of a canonical PAM, Cas12a is rapidly activated, leading to early cleavage of RPA amplicons. This depletes the template pool, limiting further amplification and reducing overall signal accumulation. In contrast, suboptimal PAMs attenuate Cas12a activation, allowing RPA to continue generating more amplicons. The resulting higher template yield leads to delayed but intensified trans-cleavage activity, ultimately producing a stronger fluorescence signal.

**Figure 3:**
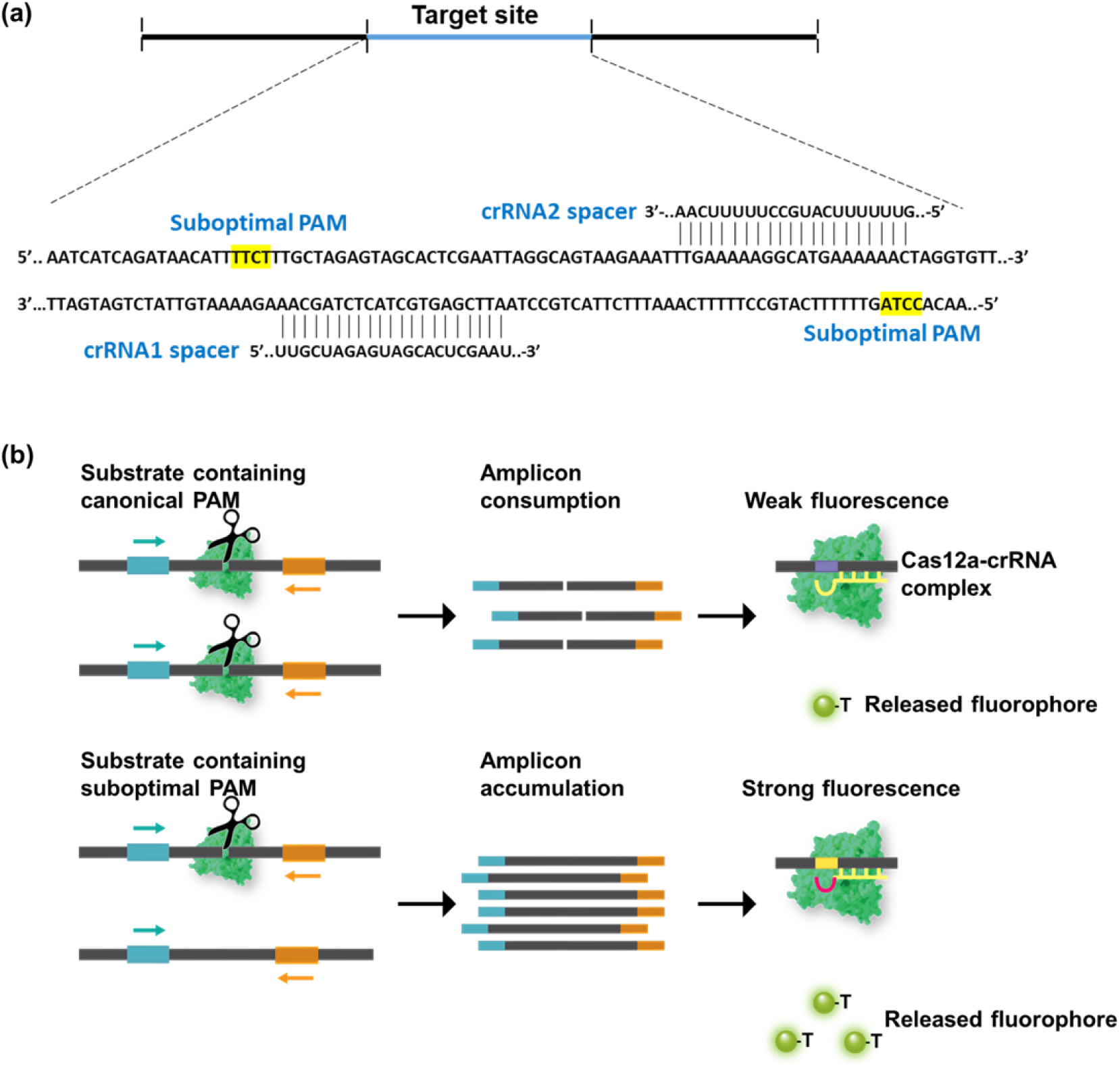
Assay design and mechanism. (a) MRSA target DNA sequence and corresponding crRNA designs featuring suboptimal PAM sequences. (b)Schematic illustration of the one-pot amplification and cleavage workflow. In the presence of a canonical PAM, Cas12a is rapidly activated, leading to early cleavage of RPA amplicons. This depletes the template pool, limits further amplification, and reduces overall signal accumulation. In contrast, suboptimal PAMs attenuate Cas12a activation, allowing RPA to continue amplifying the target sequence, resulting in greater amplicon yield and enhanced downstream fluorescence signal.

To systematically assess the contribution of each assay component and confirm that the fluorescence signal originates from the intended biochemical reaction, a panel of eight reactions (labeled #A through #H) was designed, each containing a different combination of key reagents (see table in **Fig. 4a**). The presence or absence of individual components was denoted by “+” and “-”, respectively. Among them, only the reaction #H containing all the necessary elements (including target DNA, RPA reagents, CRISPR-Cas12a complex and fluorescent probe) produced a significantly increased normalized fluorescence signal. In contrast, reactions missing one component yielded minimal to no fluorescence. These results confirm that each component is essential for assay functionality and, critically, that the fluorescence signal arises from the specific CRISPR-Cas12a-mediated trans-cleavage reaction following target recognition, rather than from any nonspecific fluorescence contributed by individual reagents. This systematic validation underscores the specificity and integrity of the one-pot assay for MRSA detection.

**Figure 4:**
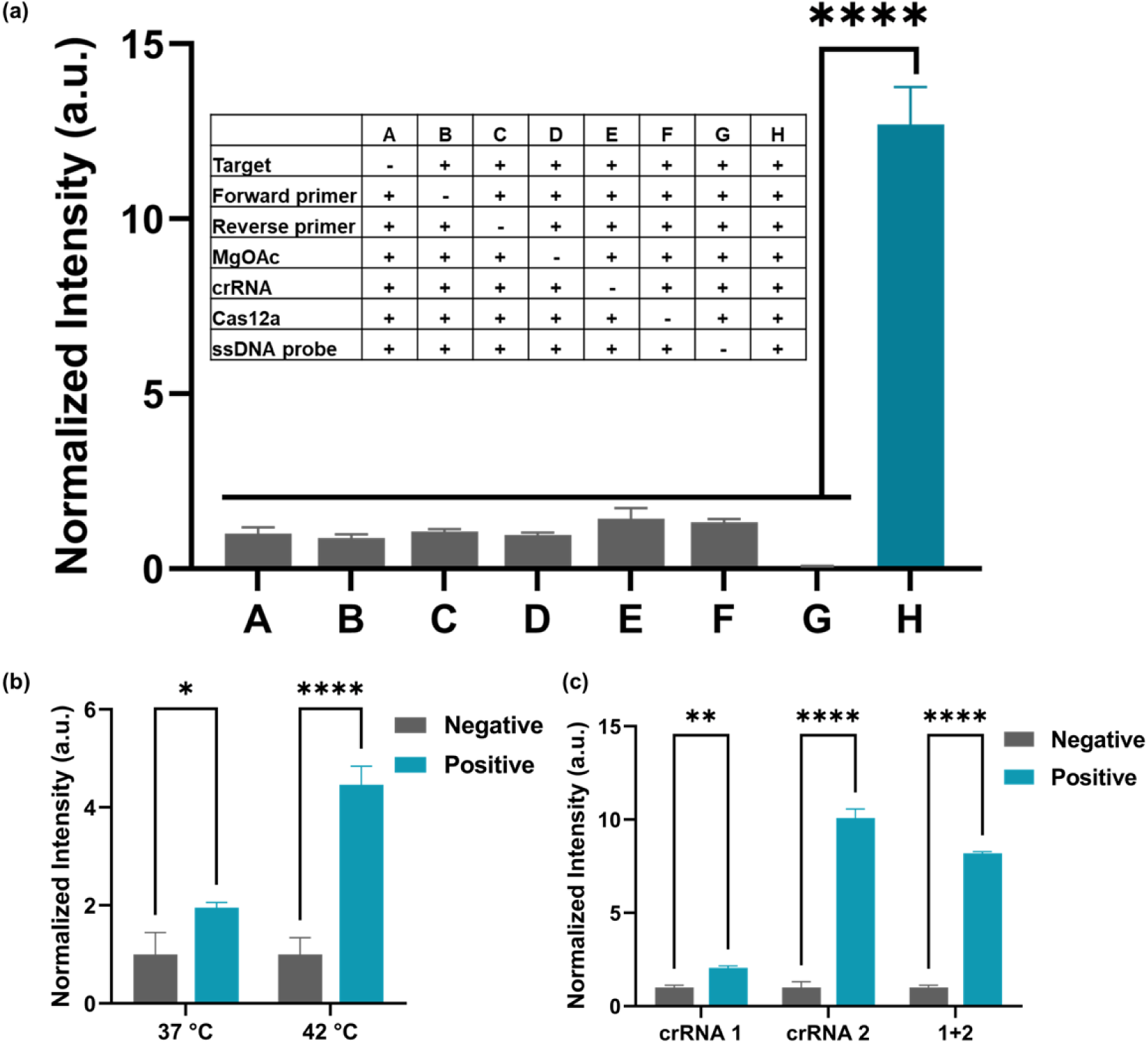
Off-chip assay evaluation. (a) Evaluation of crRNA designs for MRSA detection using fluorophore–quencher (F–Q) probes. The input target concentration is 10^7^ CFU mL^-1^. Normalized fluorescence signals are shown for each reaction. Statistical analysis was performed using one-way ANOVA. Inset: eight test reactions evaluating various combinations of assay components. Reaction #H includes all necessary components for the one-pot RPA/CRISPR-Cas12a assay, while other reactions omit specific elements to assess their contributions to signal generation. (b) Optimization of incubation temperature for the one-pot assay. The input target concentration is 10^5^ CFU mL^-1^. Statistical analysis were performed using two-way ANOVA. (c) crRNA screening for improved performance in the one-pot assay. The input target concentration is 10^6^ CFU mL^-1^ Statistical analysis were performed using one-way ANOVA. The data are represented as mean ± standard deviation (*n* = 3). For statistical analysis, *ns*, not significant = *p >* 0.05; * = 0.01 *< p* ≤ 0.05; ** = 0.001 *< p* ≤ 0.01; *** = 0.0001 *< p* ≤ 0.001; **** = *p* ≤ 0.0001.

**Fig. 4b** presents the results of temperature screening for the one-pot RPA/CRISPR-Cas12a assay. Incubation at 42 °C resulted in a significantly stronger fluorescence signal compared to 37 °C, indicating enhanced assay performance. RPA exhibits optimal activity in the range of 37–42 °C, while Cas12a typically functions best near 37 °C. At 42 °C, the elevated RPA activity promotes rapid and efficient target amplification, while the slightly reduced Cas12a activity delays trans-cleavage initiation. This temporal separation allows for sufficient amplicon accumulation prior to Cas12a-mediated probe cleavage, ultimately improving signal intensity and assay sensitivity. **Fig. 4c** evaluates the performance of two different crRNAs. Among the candidates, crRNA2, targeting a site adjacent to a suboptimal PAM, produced the highest fluorescence signal. This supports the hypothesis that reduced Cas12a activation due to suboptimal PAMs delays trans-cleavage activity, allowing more time for RPA to generate amplicons. The increased amplicon yield results in a stronger downstream fluorescence signal once Cas12a is eventually activated. crRNA1, which also targets a suboptimal PAM, generated a moderate increase in fluorescence relative to the negative control, though not as pronounced as crRNA2. Addtionaly, codelivery of crRNA1 and crRNA2 resulted in a reduced fluorescence signal compared to crRNA2 alone. Based on these results, crRNA2 was selected for subsequent experiments due to its superior signal output, optimal kinetics for delayed Cas12a activation, and its ability to reduce reagent cost by eliminating the need for crRNA combinations.

Following overnight incubation with the FAM-labeled ssDNA probe, the chip was washed four times with PBS to remove excess unbound probes. After each wash, the fluorescence of the wash solution was measured, as shown in **Fig. 5a**. A strong fluorescence signal was observed after the first wash, consistent with the high concentration of free probe applied during the labeling step. Subsequent washes led to a substantial decrease in fluorescence, and the signal plateaued after the 3*rd* and 4*th* wash, indicating the effective removal of unbound probes. This ensured that any fluorescence observed in downstream assays would originate from specific probe cleavage rather than residual unbound fluorophores. To evaluate the relationship between probe binding capacity and micropillar height, DNase was used to enzymatically cleave the immobilized ssDNA probes, and the released fluorescence was measured (**Fig. 5b**). The fluorescence intensity increased with increasing micropillar height of 100 µm, 300 µm, and 500 µm, suggesting a greater number of cleaved probes. Since the pillar diameter was kept constant, taller micropillars corresponded to higher aspect ratios, providing more surface area for probe immobilization, resulting in enhanced binding capacity. **Fig. 5c** shows the specificity of the on-chip one-pot assay using genomic DNA from three non-target bacterial strains: wild type *E. coli*, kanamycin-resistant *E. coli* DH5*α*, and wild type *S. aureus*. Bacterial DNA was extracted from bacterial cultures at a concentration of 10^8^ CFU mL^-1^. A PBS buffer solution lacking bacteria served as the no template control (NTC). The fluorescence signals generated by all bacterial samples were comparable to that of the NTC, demonstrating that the assay specifically detects the mecA gene and does not cross-react with non-target bacterial genomes. Sensitivity testing results for micropillars with heights of 100 µm and 500 µm are shown in **Fig. 5d** and **Fig. 5e**, respectively.The limit of detection (LOD) for the 100 µm pillars was 10^4^ CFU mL^-1^, while the 500 µm pillars achieved a tenfold improvement, with a LOD of 10^3^ CFU mL^-1^. This enhanced sensitivity with increased pillar height can be explained by several synergistic effects.

**Figure 5:**
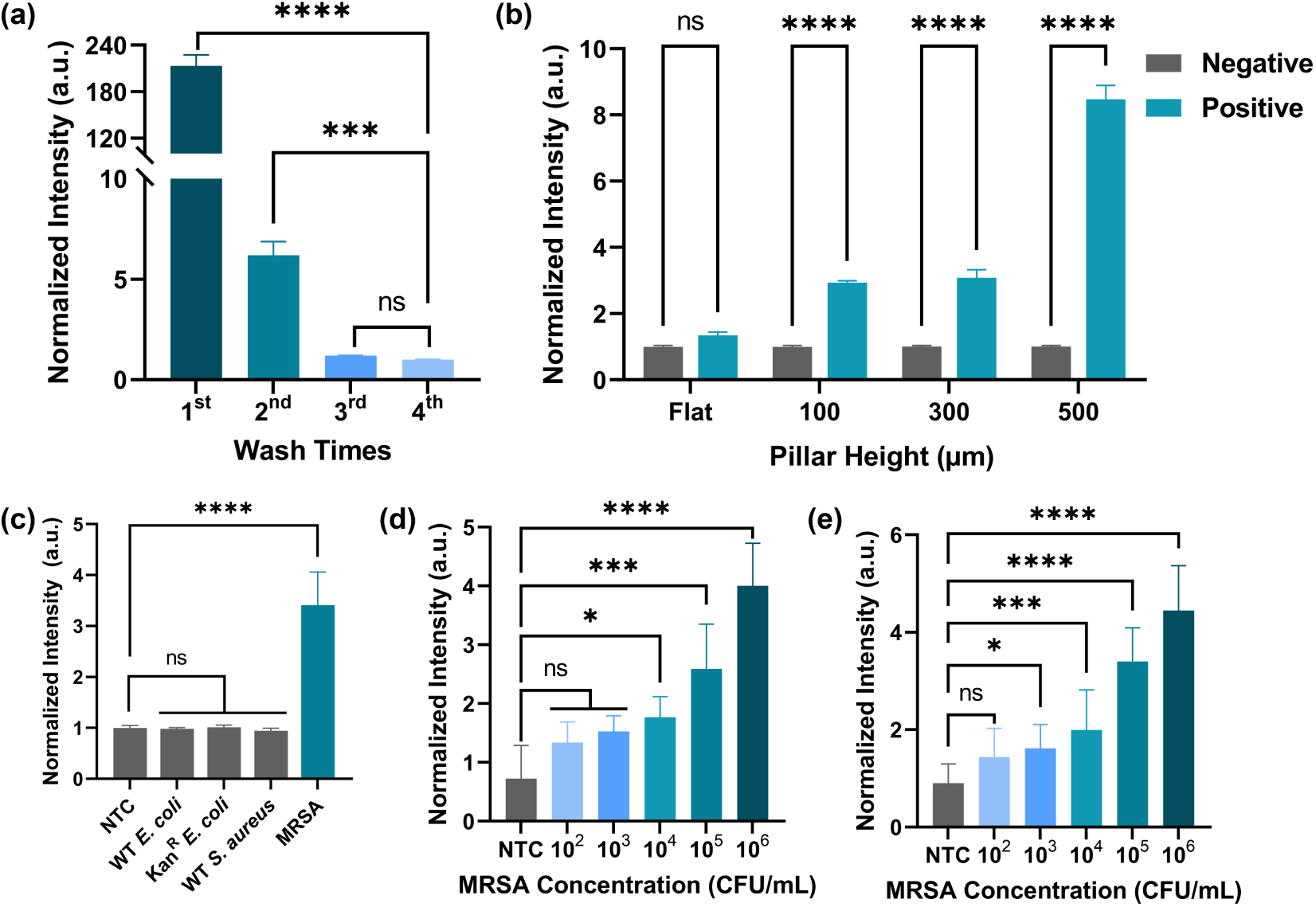
On-chip assay evaluation. (a) Comparison of normalized fluorescence intensity in the wash solution after rinsing the chip with 200 µL of PBS following probe binding. Statistical analysis was conducted using unpaired *t* -test analysis. (b) Normalized fluorescence intensity reflecting the amount of ssDNA probe bound to micropillars of varying heights. Statistical analysis were performed using two-way ANOVA. (c) On-chip specificity test using wild-type *E. coli* K12, kanamycin-resistant *E. coli* DH5*α*, and wild-type *S. aureus*, respectively. “NTC” refers to no template control. (d) Sensitivity evaluation of micropillars with a height of 100 µm. (e) Sensitivity evaluation of micropillars with a height of 500 µm. Statistical analysis were performed using one-way ANOVA. The data are represented as mean ± standard deviation (*n* = 3). For statistical analysis, *ns*, not significant = *p >* 0.05; * = 0.01 *< p* ≤ 0.05; ** = 0.001 *< p* ≤ 0.01; *** = 0.0001 *< p* ≤ 0.001; **** = *p* ≤ 0.0001.

First, taller micropillars present a larger surface area due to their higher aspect ratio, which allows for greater immobilization of ssDNA probes. A higher density of surface-bound probes increases the likelihood of interaction with activated Cas12a complexes, thereby amplifying the overall fluorescence signal. Second, the increased surface area enables higher signal accumulation per target event, which improves the signal-to-noise ratio and allows for the detection of lower concentrations of target DNA. Third, the 3D structure of taller pillars may promote more efficient diffusion and interaction of reaction components within the chip microenvironment, enhancing hybridization and cleavage kinetics. Finally, since the FAM-labeled probe lacks a quencher, background fluorescence is minimal, enabling even small changes in signal to be readily distinguished. Together, these factors contribute to the improved detection performance observed with taller micropillar structures.

We demonstrated that integrating a one-pot RPA/CRISPR-Cas12a assay with highaspect-ratio silicon micropillar arrays enhances the sensitivity of MRSA detection. By precisely fabricating micropillar structures with identical diameters but varying heights (100, 300, and 500 µm), probe binding efficiency is shown that directly scales with surface area, as validated by DNase cleavage assays. The increased three-dimensional surface provided by taller micropillars facilitated higher probe loading, resulting in improved signal output upon target recognition. Additionally, fluorescence imaging confirmed uniform probe distribution across the micropillar height, validating the effective functionalization of the entire vertical surface. These architectural advantages, combined with controlled surface chemistry (plasma treatment, APTES silanization, and glutaraldehyde coupling), established a robust and reproducible platform for surface-immobilized CRISPR detection.

When integrated with isothermal amplification like RPA, a key challenge in CRISPR-based diagnostics is the risk of premature Cas12a activation, which can result in amplicon consumption before sufficient signal accumulates. To overcome this, we designed crRNAs adjacent to suboptimal PAM sequences to delay Cas12a trans-cleavage kinetics. The crRNA screening demonstrated that suboptimal PAMs, such as TTCT, provided better assay performance than canonical sequences, likely due to reduced cleavage efficiency early in the reaction. This strategy helped temporally decouple amplification and detection phases within a one-pot reaction, maintaining simplicity while enhancing signal output. Furthermore, temperature screening indicated that 42 °C was optimal for balancing RPA efficiency with moderated Cas12a activation. These findings emphasize the importance of crRNA design and temperature tuning for optimizing one-pot CRISPR systems. Importantly, our platform achieved a detection limit of 10^3^ CFU mL^-1^ for MRSA using 500 µm micropillars—tenfold more sensitive than the 100 µm condition, while maintaining excellent specificity, as no cross-reactivity was observed with wild-type *E. coli* or *S. aureus* strains.

Despite relying on benchtop fluorescence imaging in this proof-of-concept study, the underlying design of our system is well-suited for POC diagnostics. The one-pot reaction format eliminates complex multi-step protocols and minimizes contamination risk, which are two essential features for use in decentralized or resource-limited settings. The small reagent volumes, isothermal conditions, and chip-based containment make the platform inherently portable and user-friendly. Future iterations could integrate the device with low-cost, compact detection modules such as smartphone-based fluorescence readers or colorimetric outputs, further enhancing its field deployability. While silicon substrates offer excellent reproducibility and microstructural precision, transitioning to lower-cost polymers via scalable fabrication methods (e.g., injection molding or soft lithography) could support large-scale manufacturing and commercial translation. Importantly, although this study focused on MRSA detection, the modular nature of the surface functionalization and CRISPR assay design enables easy reconfiguration for a wide range of pathogens and nucleic acid targets. In addition, the use of standard microfabrication techniques enables high-throughput production, with a single silicon wafer yielding up to 40 individual chips. This scalability supports multiplexing by enabling multiple chips, each dedicated to a distinct target, to be processed in parallel for comprehensive pathogen screening within a unified workflow. With continued development, this micropillar-enhanced CRISPR biosensor presents a promising foundation for rapid, sensitive, and accessible diagnostics at the point of care.

## 5 Conclusion

In summary, we developed a micropillar-structured silicon chip that significantly enhances CRISPR-based detection through improved probe immobilization and optimized reaction kinetics. By integrating a one-pot RPA/CRISPR-Cas12a assay with a microfabricated platform, we achieved rapid, sensitive, and specific detection of MRSA. Taller micropillars provided increased surface area, leading to greater probe density and stronger fluorescence signals, while suboptimal crRNA design enabled better control over Cas12a activation timing. The platform demonstrated tenfold sensitivity improvement and high specificity, highlighting its promise for future application in portable, contamination-free, and costeffective diagnostic systems. This work provides a foundation for further development of microstructured, CRISPR-integrated biosensors for clinical and field diagnostics

## 6 Author contributions

Conceptualization: Ruonan Peng and Ke Du. Methodology: Ruonan Peng and Ke Du. Investigation: Ruonan Peng, Zhenxuan Yuan, Demis D. John, Biljana Stamenic, Fatt Foong, Bhaskar Sharma, FNU Yuqing and Chujing Zheng. Data curation: Ruonan Peng and Ke Du. Writing-original draft: Ruonan Peng and Ke Du. Writing—review and editing: Ruonan Peng, Zhenxuan Yuan, Demis D. John, Biljana Stamenic, Fatt Foong, Bhaskar Sharma, FNU Yuqing, Chujing Zheng and Ke Du. Visualization: Ruonan Peng and Ke Du. Supervision: Ke Du. Funding acquisition: Ke Du.

## 7 Conflicts of interest

There are no conflicts to declare.

## 8 Acknowledgements

This study was supported by National Institutes of Health NIGMS R35GM142763 and NIAID R43AI179513-01.

## Notes

### Competing Interest Statement

The authors have declared no competing interest.

