## Supplement for "Silicon Micropillar-Enhanced CRISPR Biosensor for Rapid and Sensitive Detection of Drug-Resistant Bacteria"

### Experimental methods

#### Device fabrication

$< 100 >$ silicon wafers with an initial thickness of 1025 μm (University Wafer; dopant type and resistivity intentionally unspecified) were used for chip fabrication. To enable deep silicon etching, a dual-layer hardmask consisting of 150 nm of $Al_2O_3$ and a 2–3 nm $SiO_2$ capping layer was deposited using a Veeco Nexus IBD-O Ion Beam-Assisted Sputter Deposition (IBD) system. Prior to deposition, wafers were cleaned with acetone and isopropanol. Photolithography was performed using around 1.7 μm of nLOF2020 negative photoresist patterned on an I-Line Heidelberg MLA150 direct-write lithography tool. The oxide hardmask layers were subsequently etched using a $BCl_3$-based inductively coupled plasma (ICP) process (0.5 Pa chamber pressure, 500 W ICP power, 50 W RF bias, 30 sccm $BCl_3$, and 15 °C helium backside cooling). Residual photoresist was removed using sequential NMP treatment at 80 °C, followed by piranha etching and $O_2$ plasma ashing.

Deep silicon etching was carried out using a three-step Bosch process on a Plasmatherm DSE-III ICP etcher. Etch parameters were optimized via design-of-experiments (DOE) to vary aspect ratios while maintaining consistent pillar diameters. The lithographic design was fixed at 20 μm pillar diameter with 10 μm spacing. Etch depth was tuned by adjusting the dry etching time to achieve final depths of 100 μm, 300 μm, and 500 μm. Test structures were included on each wafer to characterize depth and sidewall profiles using stylus profilometry and tilted SEM imaging.

After etching, the oxide hardmasks were removed using a combination of buffered HF (BHF) and piranha solution etches. Wafers were then dry oxidized in a Tystar 8300 tube furnace to produce a conformal around 100 nm $SiO_2$ layer on all exposed silicon surfaces, passivating the micropillar structures. Finally, individual chips were singulated using an ADT 7100 dicing saw. To prevent mechanical damage to the high-aspect-ratio structures during dicing and drying, precautions were taken including reduced water flow and a final isopropyl alcohol rinse to enable gentle air-drying at room temperature.

The container was designed using SolidWorks and 3D printed with a HeyGears Relex 3D printer using Modeling PAT10 Transparent Resin. The printed part was cured in the HeyGears UltraCraft Cure system for 55 minutes at 46 °C.

The soft cap of the container was also designed in SolidWorks and fabricated using a syringe-based injection molding process. A Platinum 15A silicone mixture from BBDINO was loaded into a syringe and injected into a 3D-printed mold. The mold was then placed in an oven and cured overnight at 50 °C

#### Surface treatment

The device was immersed and incubated in 10% APTES (MP Biomedicals) dissolved in ethanol for 30 minutes with gentle shaking at 80 rpm. After incubation, it was rinsed three times with ethanol to remove excess APTES, followed by baking on a heating pad at 120 °C for 30 minutes. The device was then immersed in 2.5% glutaraldehyde (GA) solution (Thermo Scientific, prepared in PBS) and incubated for 1 hour. Afterward, it was rinsed three times with PBS to remove unbound GA and dried using an air gun. Subsequently, 20 µL of 50 µM ssDNA probe was added to the device and incubated overnight at 4 °C. The ssDNA-functionalized chip was then washed using PBS for three times, with each wash lasting 10 minutes at 80 rpm. Finally, the chip was dried with an air gun and was ready for one-pot RPA/CRISPR-Cas12a detection. Water contact angle measurements were performed using an Ossila contact angle goniometer (model L2004A-0595). Fluorescence images were captured with a Leica DMi8 fluorescence microscope equipped with a K3 digital camera and operated via Leica LAS X software.

### Bacterial culture and lysis

Methicillin-resistant *Staphylococcus aureus* (ATCC 43300) and *Staphylococcus aureus* (ATCC 25923) were purchased from Fisher Scientific. Drug-resistant *E. coli* DH5$\alpha$ (Addgene 46012, carrying plasmid pCM184 conferring resistance to ampicillin, tetracycline, and kanamycin) and *E. coli* K12 (ATCC 10798) were sourced from lab stock. All bacterial strains were cultured in tryptic soy broth (MilliporeSigma) and maintained on tryptic soy agar plates. Following overnight incubation at 37 °C with shaking at 200 rpm, 1 mL of culture was centrifuged at 8000 g for 5 min to pellet the cells. The pellet was resuspended in 1 mL of phosphate-buffered saline (PBS), and the washing step was repeated twice to completely remove residual media. Bacterial concentrations were determined by counting colony-forming units (CFU) on standard TSA plates. The washed bacteria were subsequently diluted in PBS. For DNA release, MRSA cells with different concentrations were lysed by heating at 95 °C for 10 minutes.

### DNase characterization

RNase-free DNase I (1 U/µL) was purchased from Thermo Scientific. The silicon chip was manually diced into four smaller pieces prior to use. The DNase treatment followed the manufacturer's protocol. Each reaction had a total volume of 300 µL, consisting of 30 µL of 10× reaction buffer with $MgCl_2$, 30 µL of DNase I, and 240 µL of nuclease-free water. For negative group, the DNase I was replaced by nuclease-free water. The chip fragment and the prepared reaction mixture were placed in a 0.6 mL centrifuge tube and incubated at 37 °C for 1 hour. After incubation, the chip was removed, and the fluorescence of the remaining solution was measured using an Agilent BioTek Cytation 5 imaging reader (Ex/Em = 495/520 nm).

### RPA amplification and CRISPR-Cas12a detection

TwistAmp® Basic kit was purchased from TwistDx™. The RPA primers, crRNA, LbCas12a, and single strand DNA(ssDNA) probes were obtained from Integrated DNA Technologies. Detailed sequences of all synthetic oligonucleotides are listed in Table S1. RPA primers were designed using the PrimerQuest™ Tool. NEBuffer™ r2.1 was purchased from New England Biolabs. To prepare the CRISPR-Cas12a mixture, 1.5 µL of 1 µM crRNA and 3 µL of 1 µM Cas12a were combined and incubated at room temperature for 10 minutes. Subsequently, 3 µL of 10× NEBuffer r2.1 was added. For off-chip assays, 2 µL of 10 µM ssDNA-FQ probe was included in the reaction; for on-chip assays, the probe was replaced with nuclease-free water. The RPA reaction was prepared according to the manufacturer's instructions: 29.5 µL of rehydration buffer, 13.2 µL of nuclease-free water, and 2.4 µL each of 10 µM forward and reverse primers were added to the lyophilized enzyme pellet. Then, 2.5 µL of 280 mM MgOAc was added to initiate the reaction. For one-pot reactions, 17 μL of the RPA mixture was added to the prepared CRISPR-Cas12a mixture, followed by the addition of 2 μL of MRSA lysate, resulting in a final reaction volume of 30 μL. The reaction was incubated at 37 °C for 1 hour. After incubation, fluorescence was measured using an Agilent BioTek Cytation 5 imaging reader (Ex/Em = 495/520 nm).

**Table S1**
List of synthetic oligos sequence used in this study

| Name | Sequence (5′-3′) |
|---|---|
| Forward primer | AAACAAGCAATAGAATCATCAGATAACATTT |
| Reverse primer | AAGGATCTGTACTGGGTTAATCAGTATTTC |
| crRNA 1 | UAAUUUCUACUAAGUGUAGAUUUGCUAGAGUAGCACUCGAAU |
| Target 1 | AACGATCTCATCGTGAGCTTA |
| crRNA 2 | UAAUUUCUACUAAGUGUAGAUGUUUUUUCAUGCCUUUUUCAA |
| Target 2 | CAAAAAAGTACGGAAAAAGTT |
| FQ probe | /56-FAM/TTATT/3IABkFQ |
| Cy3-labeled probe | /5AmMC6/iSp18/TTATTGCGTGAAGTTATT/3Cy3Sp/ |
| FAM-labeled probe | /5AmMC6/iSp18/TTATTGCGTGAAGTTATT/36-FAM/ |
